# Multisensory attenuation of the pupil light response in autistic and non-autistic children

**DOI:** 10.1101/2025.09.18.677095

**Authors:** Chloe Brittenham, Theo Vanneau, Sophie Molholm

**Affiliations:** Cognitive Neurophysiology Laboratory, Department of Pediatrics, Albert Einstein College of Medicine, Bronx, New York 10461, USA; Department of Pediatrics, Albert Einstein College of Medicine, Bronx, New York 10461, USA; Department of Neuroscience, Albert Einstein College of Medicine, Bronx, New York 10461, USA

**Keywords:** Multisensory, Pupillometry, Pupil, Autism, Autonomic

## Abstract

Autonomic responses to sensory stimuli are altered in autism, yet little is known about how multisensory input modulates these responses. This study examined whether auditory stimuli affect the pupil light reflex (PLR), a parasympathetically driven response to light, in autistic and non-autistic children. Pupillometry was used to measure responses to visual-only (V), auditory-only (A), and audiovisual (AV) stimuli in 72 children aged 6–14 years (34 non-autistic, 38 autistic). We hypothesized that auditory input would attenuate pupil constriction in non-autistic children and that this cross-modal modulation might differ in autism, reflecting altered sensory-autonomic functioning. Across groups, results revealed a consistent pattern: auditory stimuli elicited pupil dilation, visual stimuli evoked constriction, and simultaneous audiovisual stimuli led to attenuated constriction relative to visual-only trials. This attenuation lends support to prior findings of multisensory attenuation of the PLR. Time-binned analysis revealed a group effect during the 500–1000 ms post-stimulus window: autistic children showed significantly more positive baseline-corrected pupil responses across conditions (i.e., less constriction in V/AV and greater dilation in A), suggesting group differences in the dynamic trajectory of the pupil response. Contrary to expectations, autistic and non-autistic children did not differ significantly on peak constriction or constriction latency within visual conditions. Findings support the presence of cross-modal modulation of the PLR in both autistic and non-autistic children and suggest that auditory signals influence early-stage visual-autonomic processing similarly across groups. Pupillometry may provide a promising, noninvasive tool for probing sensory-autonomic interactions in autism. Future studies with paradigms optimized for pupil measurement may reveal more nuanced group differences and clarify links to real-world sensory challenges.

## Multisensory Processing and the Autonomic Nervous System

Navigating our complex environment requires us to process inputs from multiple sensory modalities simultaneously and ready our bodies to respond appropriately. In neurotypical individuals, multisensory processing facilitates efficient perception and behavior. By combining temporally aligned cues across modalities—such as seeing a speaker’s lips move while hearing their voice—the brain can reduce ambiguity, increase perceptual salience, and accelerate responses (Molholm et al., 2004; Molholm et al., 2002; Ross et al., 2007). For instance, audiovisual signals can trigger faster reactions than unimodal ones (Hecht et al., 2008; Mégevand et al., 2013; Molholm et al., 2002; Nidiffer et al., 2016; Sperdin et al., 2009), in part through activation of subcortical and brainstem circuits involved in autonomic regulation (Critchley et al., 2003; Nieuwenhuis et al., 2011). These systems are known to modulate arousal and facilitate attention in response to salient sensory events (Aston-Jones & Cohen, 2005; Thiele & Bellgrove, 2018).

In contrast, in autism spectrum disorder (ASD), a neurodevelopmental condition characterized by differences in social communication and the presence of restricted or repetitive patterns of behavior (APA, 2013), individuals often show reduced benefits, or even adverse effects, from multisensory stimulation (Beker et al., 2018). Behavioral and neurophysiological studies reveal diminished multisensory gain and attenuated reaction time benefits in redundant cue contexts, suggesting a less efficient or less flexible integration of sensory input (Brandwein et al., 2015; Brandwein et al., 2013; Foxe & Molholm, 2009; Kwakye et al., 2011; Molholm et al., 2020). Instead of streamlining perception, converging sensory inputs may be experienced as disjointed, unpredictable, or overwhelming, contributing to the sensory sensitivities or overload frequently reported by autistic individuals (Crasta et al., 2020; Leekam et al., 2007; Millington & Simmons, 2024). These challenges can lead to difficulties with emotion regulation, avoidance behaviors, and real-world functional impairments. For instance, many autistic individuals report feeling overwhelmed in environments with high sensory demands, such as busy classrooms, supermarkets, or social gatherings, where multiple sensory inputs (e.g., lights, voices, background noise) occur simultaneously (MacLennan et al., 2022). These experiences are often described as disorganizing or distressing and can lead to avoidance behaviors, emotional dysregulation, or shutdown (MacLennan et al., 2022). However, it remains unclear what underpins this sense of overwhelm, whether it stems from differences in attentional mechanisms, specifically in inhibition of irrelevant stimuli (e.g., Murphy et al., 2014) or from a deficit in multisensory integration leading to the experience of multiple concurrent stimuli, or both.

### Autonomic Nervous System Function in Autism

Increasingly, research points to a critical role for autonomic dysregulation in shaping these sensory experiences in autism (Gomez & Flores, 2020; Schaaf et al., 2015). The autonomic nervous system (ANS), which adjusts physiological states in response to environmental demands, is often altered in autism, with studies reporting atypical baseline arousal, parasympathetic responses, and/or sympathetic reactivity associated with autism and/or autistic traits (Anderson & Colombo, 2009; Chang et al., 2012; Daluwatte et al., 2015; Daluwatte et al., 2013; Fan et al., 2009; Kim et al., 2022; Soker-Elimaliah et al., 2023). These differences may impact the ability to regulate and maintain appropriate focused attention or arousal in response to sensory change. Importantly, autonomic dysregulation may not simply co-occur with sensory processing difficulties—it may emerge earlier and contribute to them. Longitudinal studies in infants at elevated likelihood for ASD have shown atypical pupil responses and arousal patterns before the onset of core behavioral symptoms (McCormick et al., 2018; Nyström et al., 2018; Nyström et al., 2015). Conversely, difficulties processing sensory input may amplify physiological stress, creating feedback loops that reinforce sensory and behavioral challenges. These two systems, autonomic regulation and sensory integration, likely co-develop in early life and may mutually influence one another.

### Pupillometry as a Window into Sensory-Autonomic Interactions

Pupillometry, the measurement of pupil size change, provides a promising avenue for examining the interplay between autonomic and sensory systems. The pupil’s response to light, rapid constriction followed by slower redilation, is primarily parasympathetically mediated, though redilation also involves sympathetic activation (Hall & Chilcott, 2018). This response is largely automatic but can be modulated by cortical and subcortical regions including the superior colliculus (SC) and frontal eye fields (Ebitz & Moore, 2017; Peinkhofer et al., 2019; Wang & Munoz, 2024). In contrast, the pupil’s response to the onset of sound is dilatory (Liao et al., 2016) and shows classic features of orienting responses including habituation, dishabituation, and sensitivity to stimulus magnitude (Marois et al., 2018). This dilation is closely tied to activity in the locus coeruleus-norepinephrine (LC-NE) system, a core arousal network that modulates attention and cognitive effort (Aston-Jones & Cohen, 2005). Electrophysiological and fMRI studies have demonstrated strong coupling between LC activation and pupil dynamics, with noradrenaline release accompanying dilation and simultaneously modulating auditory cortical processing (Hu & Vetter, 2024). Importantly, auditory properties such as salience can engage higher-order networks, thereby modulating arousal and driving pupil changes (Hu & Vetter, 2024). Other structures, such as the SC, also influence pupil dilation through connections to both sympathetic and parasympathetic pathways and play a key role in orienting responses to salient auditory stimuli (Wang & Munoz, 2024). These interactions underscore the integrative function of the pupil as a peripheral readout of complex sensory-autonomic dynamics.

Together, luminance- and sound-evoked pupil responses provide a physiological index of multisensory processing and the dynamic balance of the ANS. Past work in neurotypical adults has indicated that this integration of multisensory signals is additive (Liu et al., 2024; Van der Stoep et al., 2021), which may allow for the some rough inference of the relative contributions of parasympathetic and sympathetic activity. For instance, auditory input can modulate the pupil light reflex (PLR); when sound is presented simultaneously with light, the PLR is attenuated (Yuan et al., 2021), which may reflect active inhibition of parasympathetic-driven pupil constriction by concurrent auditory input, reflecting early-stage cross-modal modulation within midbrain structures such as the superior colliculus (Wang & Munoz, 2014).

### Atypical Pupil Responses in Autism

Pupillometry studies in ASD have revealed alterations in both light-evoked and sound-evoked pupil responses, suggesting disruptions in autonomic and sensory regulation. Specifically, infants with an elevated likelihood of developing ASD (by virtue of having an older sibling with ASD) show a stronger PLR (i.e., greater relative constriction amplitude; Nyström et al., 2015), and the degree of enhancement in the response was associated with later increased autistic symptomology at 3 years old (Nyström et al., 2018). However, by childhood, this relationship appears to reverse; constriction amplitude is negatively associated with greater autism symptomology (Daluwatte et al., 2015) and autistic children show longer PLR latency, smaller constriction amplitude, and lower constriction velocity as compared to their neurotypical peers (Daluwatte et al., 2015; Daluwatte et al., 2013; Fan et al., 2009). Adolescents with ASD continue to show greater constriction latency (Lynch et al., 2018) and in adulthood, increased levels of autistic traits such as restricted and repetitive behaviors have been associated with weaker and slower PLR in the general population (Soker-Elimaliah et al., 2023).

Pupil dilation in response to sound (PDR) in autism has also shown atypical patterns. For instance, infants who later develop ASD show a larger pupil response to nonsocial sounds as compared to controls (Rudling et al., 2022) and autistic children show less habituation to deviant sounds, and a larger response to aversive sounds (Song et al., 2024). These atypical pupil dynamics are consistent with findings of heightened skin conductance responses to auditory tones in autistic children (Chang et al., 2012), reinforcing theories of disruptions in the autonomic regulation in response to sound in certain contexts (Keith et al., 2019).

### The Current Study

The current study aimed to investigate whether auditory stimuli modulate the PLR differently in autistic and non-autistic children, thereby probing early-stage sensory-autonomic interactions. Using pupillometry, we measured responses to visual-only (light), auditory-only (sound), and audiovisual (simultaneous light and sound) stimuli in children between six and 14 years of age. This age range was selected because prior work suggests that multisensory processing differences are particularly prominent in childhood and early adolescence (see Crosse et al., 2022).

Although the PLR has been studied in isolation in ASD, it remains unknown how it may be modulated by auditory input. In non-autistic individuals, sounds can modulate the pupil’s response to luminance shifts when presented concurrently with luminance changes (light or dark, —attenuating constriction to light or enhancing dilation to dark (Wang & Munoz, 2014; Yuan et al., 2021)—a phenomenon proposed to reflect early multisensory integration in midbrain structures (Wang & Munoz, 2015; Wang & Munoz, 2014). If this auditory modulation of the PLR is altered in autism, it could provide a unique physiological marker of differences in early-stage multisensory integration and its autonomic correlates.

We hypothesized that, in the visual-only condition, autistic children would exhibit a weaker PLR—that is, smaller peak constriction and/or delayed constriction latency—relative to their non-autistic peers, consistent with previous reports of reduced parasympathetic response in ASD (Daluwatte et al., 2015; Daluwatte et al., 2013; Fan et al., 2009). In the auditory-only condition, we expected to observe greater pupil dilation in the ASD group, reflecting greater sympathetic activation (i.e., Chang et al., 2012). Finally, for the audiovisual condition, we hypothesized that the concurrent auditory stimulus would attenuate the PLR in the non-autistic group, as seen in previous studies of cross-modal modulation (e.g., Yuan et al., 2021), and that the degree of modulation would be altered in the autistic group, which might suggest differences in early-stage multisensory integration and/or autonomic coordination (see Beker et al., 2018).

By measuring pupil responses to light, sound, and their combination, we can assess both the strength of unimodal sensory responses and the degree to which cross-modal input shapes autonomic output. This approach has the potential to offer new insight into how autistic individuals integrate sensory information and regulate arousal, and how disruptions to these systems may contribute to everyday sensory challenges.

## Method

### Participants

The initial sample included 38 non-autistic and 54 autistic children. Following preprocessing of the data (see details in Data Preprocessing), the final sample included 34 non-autistic and 38 autistic children between the ages of six and 14 (*M*_age_ = 10.63 years, *SD* = 1.66). A licensed clinical psychologist confirmed ASD diagnoses using the Autism Diagnostic Observation Schedule 2 (ADOS-2; Lord et al., 2012) and the diagnostic criteria for autistic disorder (Diagnostic and statistical manual of mental disorders, fifth edition; DSM-5; American Psychiatric Association, 2013). A subset of the final sample (*n* = 2) were unable to complete the ADOS-2 evaluation due to masking precautions during the COVID-19 pandemic and thus the Autism Diagnostic Interview-Revised (ADI-R; Rutter et al., 2003) and the Childhood Autism Rating Scale (CARS-2; Schopler et al., 2010) were completed. Additionally, the Social Responsiveness Scale, Second Edition (SRS-2; Constantino & Gruber, 2012) was administered as a continuous measure autistic traits.

For inclusion in the non-autistic group, participants were required to have no history of neurological, developmental, or psychiatric conditions, no first-degree relatives with an ASD diagnosis, and be enrolled in a grade level appropriate for their age. Exclusion criteria for both groups included: (1) the presence of a known genetic syndrome, including syndromic ASD, (2) a history of seizure medication use within the past two years, (3) significant physical impairments, such as vision or hearing deficits, confirmed during screening on the test day, (4) preterm birth before 35 weeks gestation or a history of prenatal/perinatal complications, and (5) a Full Scale IQ (FS-IQ) below 80.

All procedures were approved by the Institutional Review Board at the Albert Einstein College of Medicine and adhered to the ethical guidelines outlined in the Declaration of Helsinki. All participants assented to the procedures and parents/guardians provided informed consent. Participants were compensated $15 per hour for their time.

**Table 1.** Participant Characteristics.

|  | Non-autistic<br>(*n* = 34) | Autistic<br>(*n* = 38) | Test statistic | *p*-Value |
| --- | --- | --- | --- | --- |
| **Age (*M*[*SD*])** | 10.36 (1.79) | 10.86 (1.52) | *t*(65.22) = -1.25 | .215† |
| **Sex (% female)** | 47.1% | 21.1% | $\chi^2(1) = 5.461$ | .019* |
| **FSIQ (*M*[*SD*])** | 104.83 (15.04) | 96.10 (15.60) | *t*(60) = 2.25 | .014* |
| **VABS-II** | - | - | - | - |
| **ABC (*M*[*SD*])** | 107.13 (10.82) | 80.93 (15.74) | *t*(59) = 7.60 | < .001*** |
| **Socialization (*M*[*SD*])** | 104.42 (10.48) | 75.80 (18.91) | *t*(44.94) = 7.28 | < .001***† |
| **Communication (*M*[*SD*])** | 108.90 (9.47) | 83.40 (15.66) | *t*(59) = 7.72 | < .001*** |
| **Daily Living Skills (*M*[*SD*])** | 105.74 (11.12) | 86.47 (17.76) | *t*(59) = 5.10 | < .001*** |
| **ADOS (*M*[*SD*])** | - | 10.62 (1.66) | - | - |
| **SRS-2 Total T score (*M*[*SD*])†** | 46.78 (6.49) | 71.38 (11.28) | *t*(47.92) = -10.53 | < .001*** |

**Note:** † *Levene's test indicated a violation of the assumption of homogeneity of variance; therefore, Welch's t-test was used for unequal variances.* Abbreviations: FSIQ, full-scale intelligence quotient; VABS-II, Vineland Adaptive Behavior Scale, Second Edition; ABC; Adaptive Behavior Composite; ADOS, Autism Diagnostic Observation Schedule; SRS-2, Social Responsiveness Scale, Second Edition.
\**p* < .05 \*\*\* *p* < .001

### Procedure

Participants were seated 70 cm from a visual display (Dell UltraSharp 1704FPT). Stimuli were presented and controlled using Presentation software (Neurobehavioral Systems) and included three conditions: a red disc (Visual; V), a 1000 Hz tone (Audio; A), and their simultaneous presentation (Audiovisual; AV). To maintain attentional engagement, participants were instructed to respond as quickly as possible by pressing a button upon detecting any stimulus. Participants’ responses were recorded using a response pad (Logitech Wingman Precision Gamepad). Stimulus onset markers were transmitted from the acquisition computer via Presentation software to log stimulus timing. Pupil size and gaze position were recorded using an SR Research EyeLink eye-tracker at a sampling rate of 500 Hz. These data were acquired under a paradigm optimized for electrophysiology recordings.

The task comprised 400 trials across four blocks, with each block containing 100 trials lasting approximately 3 minutes and 40 seconds. Each trial featured a pseudo-randomly selected stimulus (A, V, or AV), with equal probability of occurrence. The auditory stimulus consisted of a 1000 Hz tone with a duration of 60 ms, presented binaurally at 75 dB SPL. The visual stimulus was a red disc of 1.5 degrees in visual angle, positioned above a fixation cross. The audiovisual condition involved the concurrent presentation of both stimulus types (auditory and visual). Auditory stimuli were delivered through headphones, and visual stimuli were displayed on a flat-panel LCD screen. To minimize predictability, interstimulus intervals were jittered between 1000 and 3000 ms. See Figure 1A.

**Figure 1.**
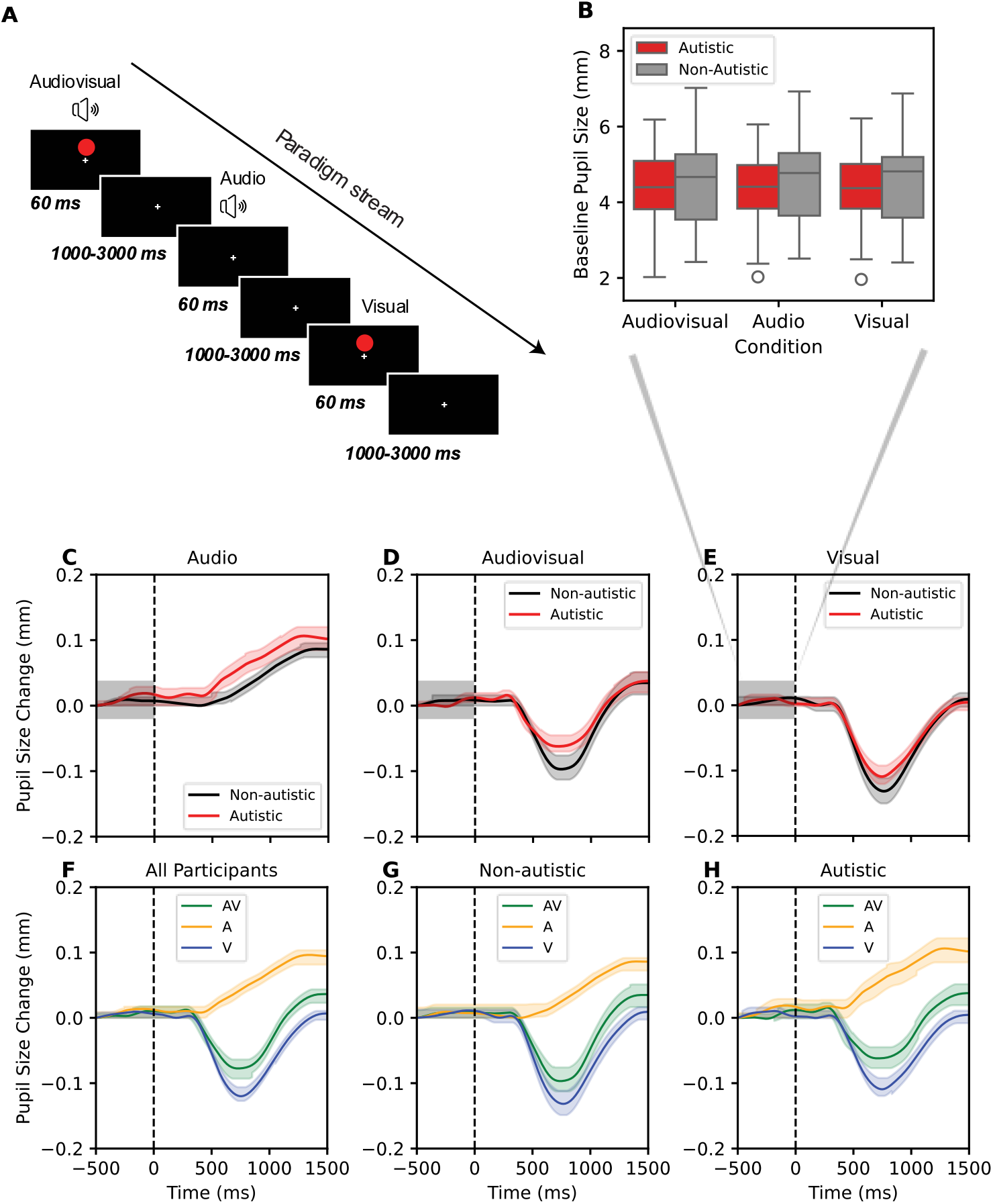
Top: (**A**) Schematic of paradigm. (**B**) Baseline pupil size across conditions, separated by group (non-autistic vs. autistic). Boxplots display the median (central line), interquartile range (box), and the full range excluding outliers (whiskers extend to 1.5 × IQR). Individual points outside this range are plotted as outliers. Bottom: Pupil size change across time for the (**C**) audio, (**D**) audiovisual, and (**E**) visual conditions, and across all conditions (AV: Audiovisual, A: Audio, V: Visual) for the (**F**) full sample, (**G**) non-autistic, and (**H**) autistic group. Dashed vertical line indicates time of stimulus onset. Shaded gray area shows baseline.

### Data Preprocessing

Data from four autistic children and one non-autistic child were excluded from preprocessing due to missing or corrupted eye-tracking files, or because the available files contained substantially reduced data resulting from poor track quality (e.g., glasses interference, eyelid occlusion). Data preprocessing was conducted using Python (v3.8) with the MNE-Python library. Raw data were epoched from -500 to 1500 ms relative to stimulus onset. Epochs containing any blinks were excluded. Offscreen data were removed, and the resulting missing data was then linearly interpolated. A 3rd-order Butterworth filter with a 4 Hz cutoff frequency was applied to smooth the pupil data. Pupil data were rescaled from arbitrary units (pixels) to millimeters to standardize the amplitude for further analysis.

Baseline correction was applied using the PLR class from the pyplr library (Martin et al., 2022). The baseline was calculated as the average pupil size before stimulus onset, and it was subtracted from all time points to obtain baseline-corrected data. Participants were excluded if their average baseline was greater than 8.1 mm or less than 1.9 mm (one non-autistic, three autistic).

Preprocessed and averaged waveforms were analyzed by condition (V, A, and AV). We applied a data-driven threshold to exclude participants contributing too few trials. The number of trials per participant per condition varied substantially (range: 1–119 trials, *M* = 40.7, *SD* = 30.9), with a positively skewed distribution (skewness = 0.73, *SE* = 0.155). To address this variability while retaining a representative sample, participants’ conditions where they contributed fewer than 12 trials (the 25th percentile) were excluded from analyses. In some cases, this resulted in all conditions being excluded for a particular participant. Within the non-autistic group, 79.4% of participants contributed data from all three conditions, 11.8% had data from two conditions, and 8.8% from one condition. Within the autistic group, 89.4% of participants contributed data from all conditions, 5.3% from two conditions, and 5.3% from one condition.

Of the 86 participants who were initially included in the preprocessed dataset, ten (two non-autistic, eight autistic) were excluded due to insufficient trials and four (one non-autistic, three autistic) were excluded for exceeding the limits of expected baseline pupil, resulting in the final sample size of 72. An average of 49 (*SD* = 29) trials per participant, per condition, were retained after cleaning and exclusion (A: *M* = 47, *SD* = 27; AV: *M* = 47, *SD* = 29, V: *M* = 52, *SD* = 32).

### Statistical Analysis

The following pupil measures were computed for all conditions: baseline pupil size, average pupil size change, average pupil size change binned, and constriction and dilation metrics. For the V and AV conditions, pupil constriction metrics were derived using the PyPlr library (Martin et al., 2022). Baseline pupil size was defined as the average pupil diameter during the pre-stimulus period (–500 to 0 ms). Average pupil size change was calculated as the mean deviation from baseline across the full epoch, as well as within specific time windows (0–500 ms, 500–1000 ms, and 1000–1500 ms) to capture temporal dynamics. Pupil dilation was indexed by the maximum pupil size increase from baseline within the epoch, and dilation latency was defined as the time at which this maximum occurred. Pupil constriction metrics included constriction latency (the time from stimulus onset to the beginning of constriction), peak constriction (the smallest pupil size following stimulus onset), constriction amplitude (the difference between baseline and peak constriction), and peak constriction latency (the time at which peak constriction occurred).

Preliminary analyses examined potential age effects and trial loss rates. There were no associations between any of the pupil metrics and age (*p*s > .20), thus age was not included in the subsequent models. Significantly more trials were retained in the A and AV conditions as compared to the V conditions (*p*s < .001), however, trial loss did not differ between groups (*p* = .101) and there was no interaction between group and condition for number of retained trials (*p* = .538).

We also tested whether missingness in condition-level data was systematically associated with diagnostic group. Chi-square tests indicated that the likelihood of missing data in any given condition (A, AV, or V) did not differ significantly between autistic and non-autistic groups (all *p*s > .31), supporting the assumption that data were missing at random (MAR). Therefore, all data that passed preprocessing and trial count thresholds were included in the subsequent analyses. Linear mixed-effects models (LMMs) were selected, as they can accommodate partially missing within-subject data under MAR. All pupil measures were analyzed using LMMs, with group and condition as fixed effects and participant as a random effect to account for individual differences and within-subject dependency.

All statistical analyses were conducted using Jamovi (The jamovi project, version 2.3.28, https://www.jamovi.org) or SPSS (IBM Corp., Armonk, NY; version 30). Pupil waveforms plotted by group and condition are shown in Figure 1C–H.

## Results

### Baseline Pupil Diameter

Baseline pupil was calculated as the average pupil size between the start of the epoch and the onset of stimulus (V: light, A: sound, or AV: light and sound). LMM was conducted with group (non-autistic, autistic) and all conditions (A, AV, V) as fixed factors and participant as a random effect. There were no significant main effects or interaction effect (*p*s > .20) for baseline pupil diameter. See Figure 1B.

### Average Pupil Size Change (Full Trial)

Average pupil size change was computed as the mean deviation from baseline across all time points in each epoch. LMM was conducted with group (non-autistic, autistic) and condition (A, AV, V) as fixed factors and participant as a random effect. There was a main effect of condition, *F*(2, 126.1) = 40.29, *p* < .001, with post hoc pairwise comparisons showing significant differences between A and AV, *t*(140) = 6.16, *p* < .001 and A and V, *t*(139) = 8.58, *p* < .001, and a marginal difference between AV and V (*p* = .058), see Figure 2B. Relative to baseline, pupil size increased (dilated) in the A condition, decreased (constricted) in the AV condition, and showed the greatest constriction in the V condition. There was no main effect of group (*p* = .134) and no significant group by condition interaction (*p* = .681).

**Figure 2.**
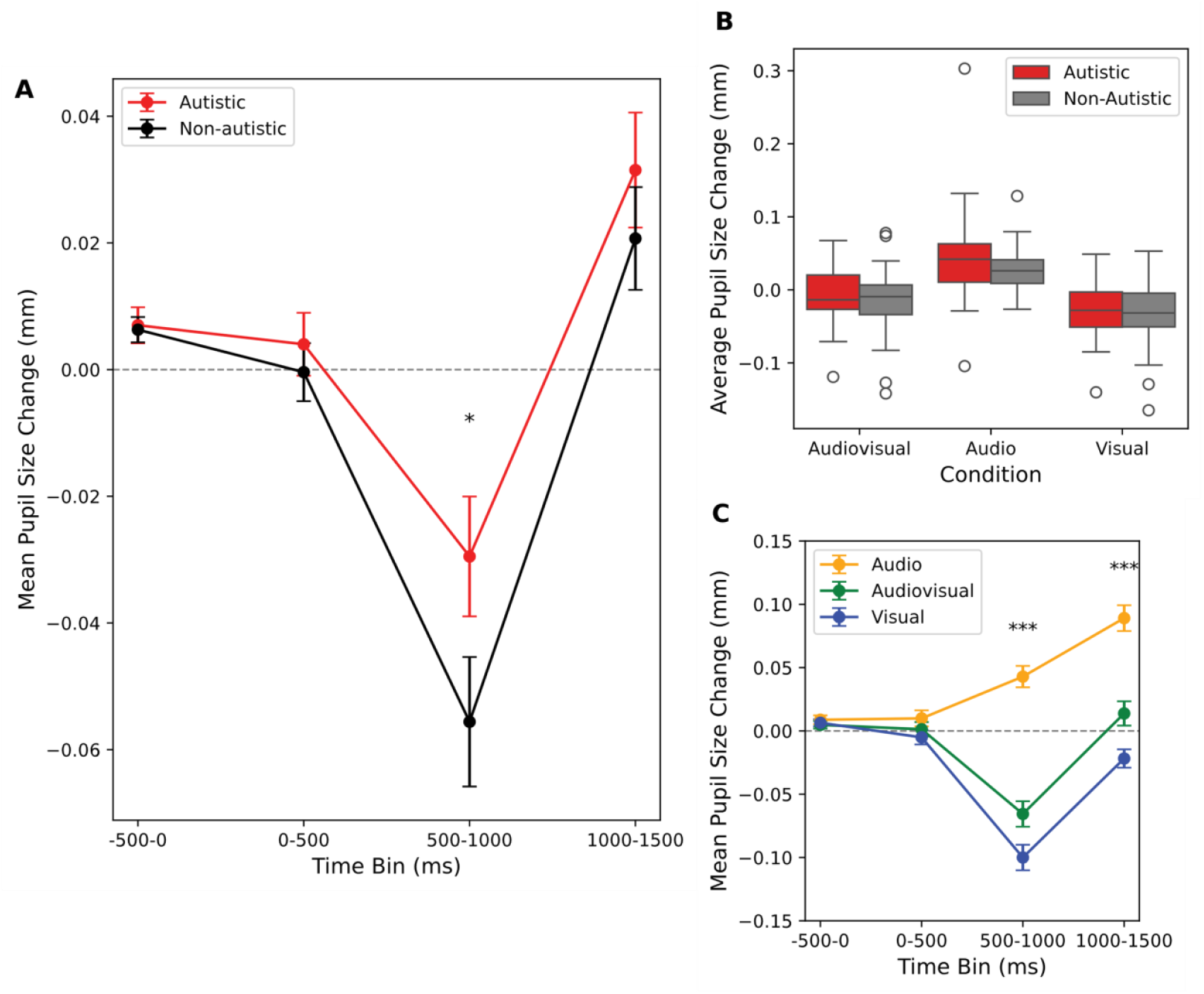
(**A**) Mean pupil size change per bin for the two groups (collapsed across conditions). (**B**) Average pupil size change. Boxplots display the median (central line), interquartile range (box), and the full range excluding outliers (whiskers extend to 1.5 x IQR). Individual points outside this range are plotted as outliers. (**C**) Mean pupil size change per bin across the three conditions (collapsed across groups). * *p* < .05 *** *p* < .001

**Figure 3.**
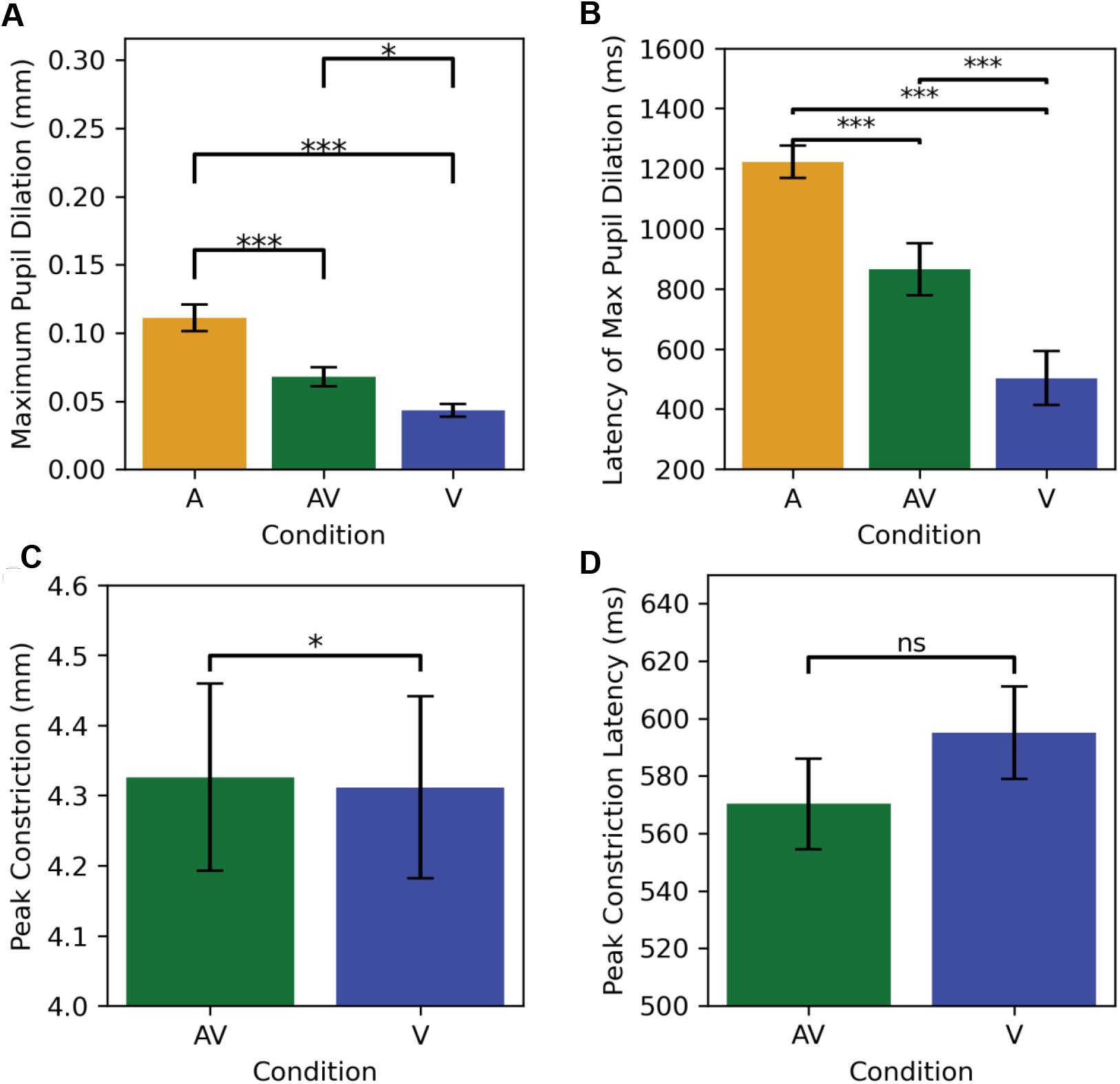
(**A**) Average maximum pupil dilation (mm) and (**B**) latency of maximum pupil dilation (ms) for the three conditions, A: Audio, AV: Audiovisual, V: Visual. (**C**) Peak constriction pupil size in mm and (**D**) latency of peak constriction (ms) for the visual conditions. Error bars represent 1 SE above and below the mean. * *p* < .05 *** *p* < .001 ns = non-significant.

### Average Pupil Size Change (Binned)

Average pupil size was computed by bin, relative to stimulus onset at time 0 for –500–0 ms, 0–500ms, 500–1000ms, 1000–1500ms. LMMs were run for each bin separately with group (non-autistic, autistic) and condition (A, AV, V) as fixed factors and participant as a random effect. The first two time bins (–500–0 ms, 0–500ms) showed no significant effects (*p*s > .17), however, significant effects were identified for the last two time bins (500–1000 ms, 1000–1500 ms).

LMM for the 500–1000 ms bin showed significant effects of condition, *F*(2, 132.49) = 63.11, *p* < .001 (see Figure 2C), and group, *F*(1, 66.06) = 5.13, *p* = .027 (see Figure 2A). Follow-up pairwise comparisons showed significant differences between each condition (A > V: *t*(139) = 10.62, *p* < .001; A > AV: *t*(140) = 8.02, *p* < .001; AV > V: *t*(141) = 2.53, *p* = .013), with dilation in the auditory condition (*M* = .04, *SE* = .009), constriction in the AV condition (*M* = -.067, *SE* = .009) and greater constriction in the V condition (*M* = -.101, *SE* = .009). The non-autistic group (*M* = -.054, *SE* = .008) showed significantly smaller overall baseline-corrected pupil size compared to the autistic group (*M* = -.029, *SE* = .007). There was no group by condition interaction (*p* = .894).

LMM for the 1000–1500 ms bin also showed a significant effect of condition, *F*(2, 126.83) = 47.44, *p* < .001, but no group effect or interaction effect (*p*s > .35). Follow up pairwise comparisons showed significant differences between all conditions, as in the 500–1000 ms bin (A vs. V: *t*(138) = 9.41, *p* < .001; A vs. AV: *t*(139) = 6.31, *p* < .001; V vs. AV: *t*(139) = 3.03, *p* = .003).

### Pupil Dilation

Maximum pupil size was computed as the maximum increase from baseline during the epoch and maximum pupil size latency was the time of maximum pupil size. LMM was conducted for maximum pupil dilation with group (non-autistic, autistic) and all conditions (A, AV, V) as fixed factors and participant as a random effect. A significant effect of condition, *F*(2, 128.4) = 30.53, *p* < .001 was found. Follow up tests showed significant differences between each condition (A vs. V: *t*(137) = 7.61, *p* < .001; AV vs. V: *t*(138) = 2.79, *p* = .018; AV vs. A: *t*(137) = 4.77, *p* < .001), with the greatest maximum dilation in the A condition (*M* = .109 mm, *SE* = .007), then the AV condition (*M* = .068 mm, *SE* = .007), and then the V condition (*M* = .045 mm, *SE* = .007). No effect of group or interaction was found (*p*s > .20).

LMM for maximum pupil dilation latency with group (non-autistic, autistic) and all conditions (A, AV, V) as fixed factors and participant as a random effect showed a significant effect of condition, *F*(2, 131.27) = 26.42, *p* < .001. Follow up tests showed significant differences between each condition (A vs. V: *t*(138) = 7.15, *p* < .001; AV vs. V: *t*(139) = 3.49, *p* = .002; AV vs. A: *t*(139) = 3.62, *p* = .001), with maximum dilation achieved in the V condition first (*M* = 511 ms, *SE* = 76.9), then the AV condition (*M* = 855 ms, *SE* = 78.1) and then the A condition (*M* = 1215 ms, *SE* = 78.1). No significant interaction or main effect of group was found (*p*s > .55).

### Pupil Constriction

Pupil constriction was assessed for the V and AV conditions. The following measures of constriction were obtained using the Pyplr library (Martin et al., 2022): constriction latency (i.e., time from stimulus onset to the beginning of constriction), peak constriction (i.e., the smallest pupil size, in mm), constriction amplitude (i.e., the absolute difference between baseline and peak constriction), and peak constriction latency (i.e., the index at which peak constriction occurred).

LMMs were conducted separately for constriction latency, peak constriction, constriction amplitude, and peak constriction latency with group (non-autistic, autistic) and visual conditions (V, AV) as fixed factors and participant as a random effect. For peak constriction, there was a main effect of condition, *F*(1, 63.1) = 4.64, *p* = .035, where the simultaneously presented auditory stimulus significantly attenuated the peak constriction of the pupil in response to light. No main effect of group (*p* = .208) or interaction effect (*p* = .756) was found. No main effects or interaction effects were found constriction amplitude, constriction latency, or peak constriction latency (*p*s > .06).

## Discussion

The pupil light reflex (PLR), the rapid constriction of the pupil in response to light, is mediated by the parasympathetic pathway and provides a sensitive index of autonomic function. In autism, prior studies have reported alterations in the PLR, including reduced constriction amplitude and delayed constriction latency, suggesting differences in autonomic regulation (Daluwatte et al., 2015; Daluwatte et al., 2013; Fan et al., 2009). In parallel, a growing body of research has demonstrated that multisensory processing, the ability to integrate information across sensory modalities, is atypical in autistic children, particularly during middle childhood and adolescence, when these systems are still undergoing developmental refinement (Beker et al., 2018; Crosse et al., 2022; Foxe et al., 2015), suggesting that both sensory and autonomic systems may follow altered developmental trajectories in autism. Yet little is known about how these systems interact, for instance, whether auditory input modulates visual-autonomic responses differently in autism. The current study examined pupil responses to light, sound, and their combination in autistic and non-autistic children, with a particular focus on whether auditory stimuli modulate the pupil’s response to light differently across groups.

Contrary to prior research demonstrating a weaker or delayed PLR in autistic children (Daluwatte et al., 2015; Daluwatte et al., 2013; Fan et al., 2009), we found no significant group differences in peak constriction amplitude or latency in the visual-only or audiovisual conditions. Thus, classic PLR metrics did not indicate altered parasympathetic function in autism in this sample. Across conditions, the time-binned analysis, which integrated information across time rather than relying on a single peak value, revealed a significant group effect during the 500– 1000 ms window following stimulus onset, a period encompassing both peak constriction, in the visual and audiovisual conditions, and initial dilation response, in the auditory-only condition. As indicated by the main effect of group, autistic children showed more positive pupil responses across conditions relative to non-autistic children during this window. This effect may point to subtle group-level differences in the dynamic trajectory of pupil response and be suggestive of less constriction to light (Daluwatte et al., 2015; Daluwatte et al., 2013; Fan et al., 2009), greater phasic dilation in response to the auditory component (Chang et al., 2012), or both in the autistic group. However, because this effect was present only as a main effect, and no group x condition interaction was observed, it should be interpreted as a general shift in pupil dynamics rather than a condition-specific alteration.

Notably, the lack of significant group-by-condition interaction indicates that while overall pupil dynamics differed, the way in which auditory input modulates the PLR remains similar across autistic and non-autistic children. Consistent with findings in neurotypical adults (Wang & Munoz, 2014; Yuan et al., 2021), the PLR was attenuated when the visual stimulus was presented concurrently with sound, indicating cross-modal modulation of early autonomic responses, potentially mediated by active inhibition of the parasympathetic constriction pathway (Yuan et al., 2021). This supports the idea that auditory stimuli can modulate visual reflexes through mechanisms such as early sensory integration (Wang & Munoz, 2015; Wang & Munoz, 2014). The pattern of results—greater pupil constriction in the visual condition, intermediate constriction in the audiovisual condition, and dilation in the auditory-only condition—suggests an additive interplay between parasympathetic and sympathetic systems (Liu et al., 2024; Van der Stoep et al., 2021). Our results have expanded these findings to both a pediatric sample, and a clinical sample.

## Limitations

A limitation of the current study is that the paradigm was not originally optimized for extracting classic PLR metrics, but instead for EEG recordings. As a result, stimulus duration and trial timing potentially introduced carryover effects or limited time for recovery. In addition, more trials were lost for the visual condition than for the auditory and audiovisual conditions, though trial loss rates were comparable across groups. It is noteworthy that despite these constraints, the number of usable trials retained per participant in our final dataset exceeded the number typically reported in prior PLR studies, where fewer than 10 trials per condition are often used (e.g., Daluwatte et al., 2015; Daluwatte et al., 2013; Fan et al., 2009; Soker-Elimaliah et al., 2023). This higher trial rate may have enhanced the reliability of our within-subject measures and improved our ability to detect subtle effects in time-binned analyses.

## Future Directions

The present findings highlight pupillometry as a promising, noninvasive tool for probing autonomic and multisensory processes in autism. Because constriction reflects parasympathetic activity and dilation reflects sympathetic arousal, pupil measures can provide a sensitive assay of autonomic balance, and they do so without requiring explicit behavioral responses (Yuan et al., 2021). This makes them particularly useful for studying individuals with higher support needs, including those who are non-speaking, for whom traditional task-based measures may not be feasible. Future work should include these types of participants, and employ paradigms optimized for pupillometry, using more intense light stimuli and longer intertrial intervals to capture the full constriction–redilation cycle. Such designs may improve sensitivity to group differences and clarify how auditory input modulates visual-autonomic responses.

Importantly, longitudinal work will be needed to map developmental trajectories, as both multisensory integration and PLR dynamics appear to follow atypical courses across infancy, childhood, and adolescence in autism (Crosse et al., 2022; Daluwatte et al., 2013; Fan et al., 2009; Nyström et al., 2015). For example, while PLR latency typically decreases with age in non-autistic children, this trajectory is absent in autism (Daluwatte et al., 2013), and early enhancements in pupil responses observed in infants later diagnosed with autism (Nyström et al., 2018) may give way to blunted responses in older children (Daluwatte et al., 2015; Fan et al., 2009). This pattern suggests that autonomic and multisensory abnormalities shift over development and that pupillometry could serve as a useful longitudinal marker of these maturational changes.

Finally, future work should aim to directly link pupil-based indices of sensory responsiveness to real-world sensory sensitivities or adaptive functioning as this could help clarify how autonomic dynamics relate to the lived experience of autistic individuals which will be essential for clarifying the clinical and translational relevance of multisensory-autonomic differences in autism.

## Data Availability

Data available upon request

## Ethical Information

All procedures were approved by the Institutional Review Board at the Albert Einstein College of Medicine (IRB approval #2021-13433) and adhered to the ethical guidelines outlined in the Declaration of Helsinki. All participants assented to the procedures and parents/guardians provided informed consent. Participants were compensated $15 per hour for their time.

